# Beyond the connectome: A map of a brain architecture derived from whole-brain volumetric reconstructions

**DOI:** 10.1101/2020.05.24.112870

**Authors:** Christopher A. Brittin, Steven J. Cook, David H. Hall, Scott W. Emmons, Netta Cohen

## Abstract

Animal nervous system organization is crucial for all body functions and its disruption can manifest in severe cognitive and behavioral impairment. This organization relies on features across scales, from nano-level localization of synapses, through multiplicities of neuronal morphologies and their contribution to circuit organization, to the high level stereotyped connections between different regions of the brain. The sheer complexity of this organ means that to date, we have yet to reconstruct and model the structure of a complete nervous system that is integrated across all these scales. Here, we present a complete structure-function model of the nematode *C. elegans* main neuropil, the nerve ring, which we derive by integrating the volumetric reconstruction from two animals with corresponding synaptic and gap junctional connectomes. Whereas previously the nerve ring was considered a densely packed tract of axons, we uncover internal organization into 5 functional bundles and show how they spatially constrain and support the synaptic connectome. We find that the *C. elegans* connectome is not invariant, but that a precisely wired core circuit is embedded in a background of variable connectivity, and propose a corresponding reference connectome for the core circuit. Using this reference, we show that the architecture of the *C. elegans* brain can be viewed as a modular Residual Network that supports sensory computation and integration, sensory-motor convergence, and brain-wide coordination. These findings point to scalable and robust features of brain organization that are likely universal across phyla.

## Introduction

A primary goal of systems neuroscience is to understand how the brain’s structure and function combine to generate behavior. Since the discovery of neurons and their connections through synapses and gap junctions, a major effort has focused on characterizing these units and the micro- and macro-circuits that they comprise, culminating in a growing body of high resolution nano-connectomic data across species^1–10^. Naturally, data, however rich, cannot, on their own provide explanatory power to address the computation within circuits or to determine how these circuits communicate and coordinate information flow to generate behavior. Indeed, constructing a comprehensive brain map will require a meaningful strategy for integrating structure and function across scales. Achieving this feat in even a small animal can provide a useful model for postulating principles of organization across scales^11^.

The free-living nematode *C. elegans* has a small, compact nervous system^2^ while exhibiting a range of complex, individualized behaviors, making it an ideal model system for studies of whole brain organization^11^. All 302 *C. elegans* neurons have been anatomically characterized based on serial sectioned electron micrographs (EM)^2^ to produce a whole animal connectome^2,5,12^. This animal’s invariant cell-lineage^13^ and anatomy^2^ might suggest that its connectome too is invariant^14^. Unfortunately, the small sample size of available reconstructions has precluded a reliable estimate of reproducibility and variability of the synaptic connectome. Furthermore, while the synaptic wiring has been exhaustively characterized^2,5,12^, the spatial proximity of neurons is only partially determined^15,16^. Thus, it remains to be determined whether lessons about whole brain organization in *C. elegans* can inform questions and approaches for other systems.

We provide two complete volumetric reconstructions of the *C. elegans* nerve ring from legacy EMs^15^, one from a young adult and one from a larval stage 4 (L4) animal (Methods, Table S1, Figure S1). Both EM series approximately span the same 36 *µ*m long volume, starting and ending in the anterior and ventral ganglion, respectively (Figure 1(a)). Our reconstructions provide a complete, nano-scale resolution dataset of all axon-axon physical contacts in the nerve rings of these two animals. We define two neurons as neighbors if their axons are physically adjacent in at least one EM section^15^. To characterize synaptic pathways within a spatial context, we integrated our volumetric reconstruction with our recent rescoring of synapses on the same L4 and adult animals^5^ (Supplementary Online Materials, SOM).

**Figure 1.**
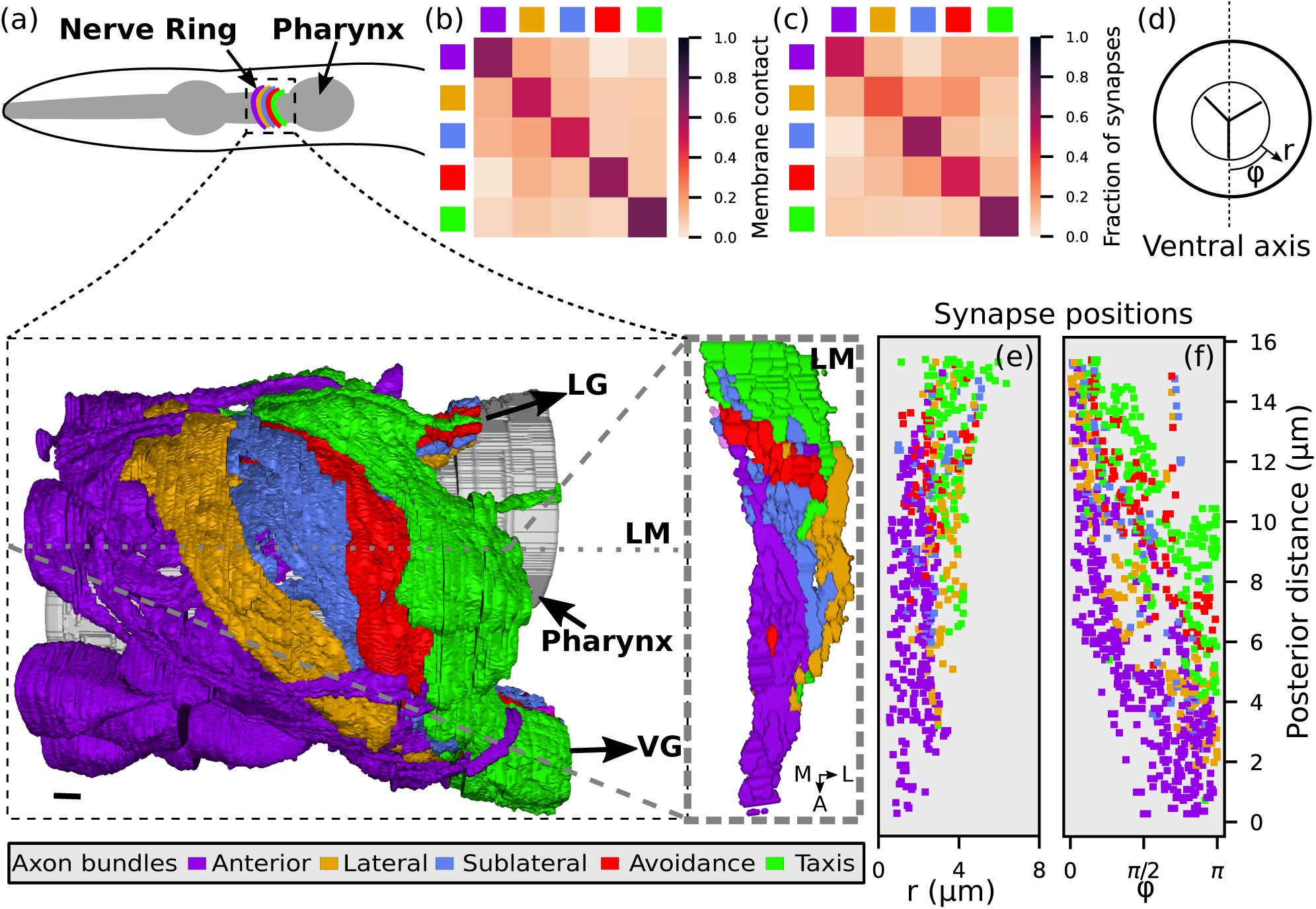
The nerve ring is organized into 5 major axon bundles that are ordered along the AP axis. (a) The nerve ring is the most synaptically dense region of the nervous system and occupies <4% of the worm’s length. 181 cells from both head and tail ganglia project axons into the nerve ring and organize into 5 bundles: Anterior, Lateral, Sublateral, Avoidance and Taxis (see text). The complete volumetric reconstruction of the 181 axons spans 36 *µ*m (Figure S1). A 15 *µ*m-wide region (inset) is shown, viewed from left, with superficial axons that obscure the AP ordering removed. A 250 nm oblique volumetric slice at approximately the lateral midline (LM) rendered with no axons removed (right). A: anterior, M: medial, L: lateral, LG: lateral ganglia, VG: ventral ganglia. Scale bar: 1 *µ*m. (b) The distribution of 𝔸^4^ intra and inter-bundle axon-axon membrane contacts. (c) The distribution of intra- and inter-bundle synapses (including 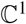 – 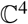 synapses with polarity from row onto column). (b,c) normalization: rows sum to 1. (d) Position in nerve ring represented in cylindrical coordinates. (e) Projection of synapse locations onto (r,z) plane. AP distance *z* measured relative to the anterior-most point of the nerve ring. (f) Projection of synapse locations onto (φ,z) plane (right side, L4 reconstruction).

### Axonal trajectories support a targeted and precise core circuit that is embedded in a variable background

Consistent with^15^, our volumetric reconstructions show that axonal trajectories are bilaterally (left/right) conserved (Supplementary Results, Figure S2). We therefore hypothesized that bilateral symmetry of *C. elegans* axons could extend to the nanoscale to support a homology of physical contacts and synapses between cells. Homologous neurons exhibit similar neighborhood sizes (Figure S3(b)) with statistically high overlaps in neighborhood composition (Figure S3(c)). However, the smallest 35% of axon contacts (<0.4*µ* m^2^) are not reproducible (Figure S4(a)), account for only 2% of total membrane contact area between all neurons (Figure S4(b)) and contain less than 10% of synaptic contacts (Figure S5(a)). As these small contacts are likely ‘spurious’, we exclude them from our analysis. We conclude that axonal trajectories are highly conserved and that these conserved trajectories are finely resolved to enforce a stereotyped pattern of cell-cell contacts.

The availability of two reconstructions, combined with the bilateral homology of the nerve ring, naturally lends itself to establishing a reference dataset that is more likely spatially conserved across animals, providing a basis to address mechanistic questions about precision and variability of the connectome at nanoscale resolution. We defined the adjacency graph, 𝔸^*δ*^, of axon contacts across 4 datasets (adult left, adult right, L4 left and L4 right datasets), where *δ* labels the conserved degree (axon contact sites in 𝔸^*δ*^ occur in 1 ≤ *δ* ≤ 4 datasets). The 𝔸^4^ consensus dataset comprises ∼40% of all axon contact sites (Figure S6(a)) and exhibits above average membrane contact area (Figure S6(b)). Adjacency graphs of chemical synapses 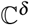 and gap junctions 𝔾^*δ*^ are similarly defined. We define 𝔸^4^, 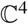 and 𝔾^4^ contacts as reference datasets and hypothesize that the 𝔸^4^ set of axon contacts is representative of the conserved physical contacts across individuals in *C. elegans* and is more likely to support a conserved synaptic connectome.

To examine this hypothesis, we exploit the combined spatial and synaptic information across datasets over the entire neuropil. We assume that stereotyped wiring patterns require precision to find target neurons and specificity to avoid off-target neurons, and formulate statistical models of axonal and synaptic connectivity to capture the relative propensity of a contact to occur in 1, 2, 3 or all 4 of the datasets (Methods). We find that a minimal model with three parameters suffices (Methods); these are the fraction of target contact sites, *f*, the precision, *p*, for target connectivity, and the frequency to avoid off-target contacts or specificity, *s*. Despite their parsimony, these models yield good fits for the relative distribution of axon contacts, synapses and gap junctions across the 4 datasets (Methods, Figure 2(a)). The high reproducibility of axon contacts across datasets (𝔸^4^ count) is consistent with our model prediction that less than half of axonal contacts made are actively targeted (*f* = 0.44, Figure 2(a)) with high precision (*p* = 0.95). The significant variability across datasets is accounted for by a non-negligible basal axon contact rate of 20-25%.

**Figure 2.**
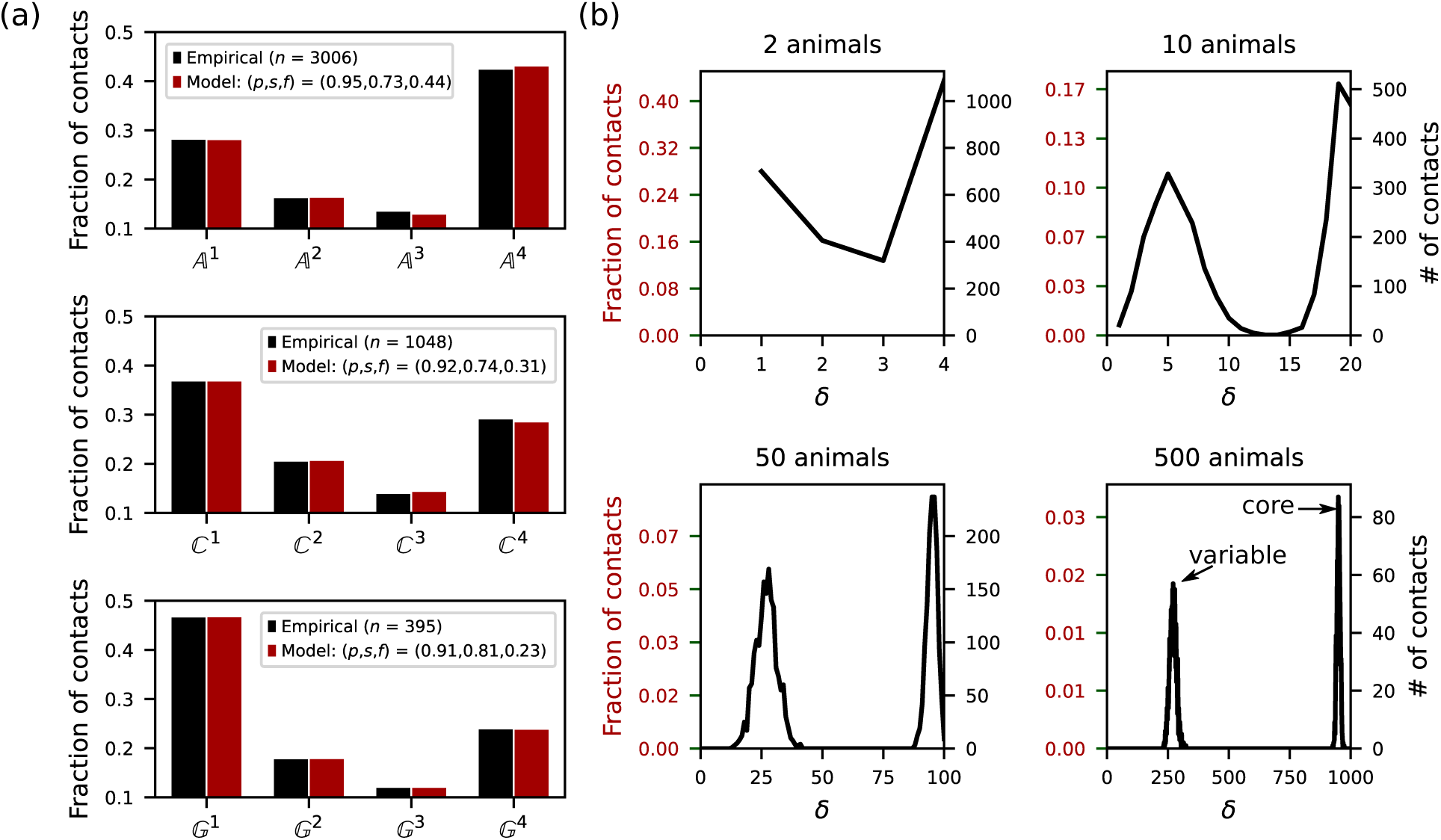
The nerve ring is comprised of a core circuit embedded in a variable background. (a) Empirical data and model fits for the reproducibility, across *δ* datasets, of axon contacts, 𝔸^*δ*^ (top), chemical synapses, 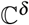 (middle) and gap junctions, 𝔾^*δ*^ (bottom). Empirical and model counts are normalized by the total empirical count of contact sites, *n* (e.g., for axon contacts, 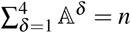). (b) Surrogate data for 4, 20, 100, 1000 datasets (obtained from 2, 10, 50 and 500 model animals). 4 datasets suffice to deduce that the distribution is bimodal (hence requiring two mechanisms) and grossly separable into the low homology (non-specific, non-target) and high homology (precisely targeted) contacts, with a split of 1 − *f*:*f*. 20 datasets (10 animals) would suffice to completely distinguish between the core and variable subcircuits. Due to finite precision, no single connection is expected to be conserved across 500 animals. In each dataset, 75% of the circuit consists of target contacts.

How useful is the 𝔸^4^ reference in predicting conserved connections? 86% of the 𝔸^4^ contact sites and 56% of the 𝔸^3^ contact sites constitute the vast majority of the predicted core circuit (Methods). Larger membrane contacts make up more than 80% of 𝔸^4^ contact sites and are more reproducible (the higher precision, *p* = 0.98, and larger fraction, *f* = 0.76, indicate that 96% of larger contacts are likely conserved, Figure S7). We conclude that the 𝔸^4^ dataset offers an excellent candidate set of conserved axon contacts. We further estimate that 20 datasets (from 10 animals, with each bilateral reconstruction yielding 2 datasets) would be sufficient to identify all contact sites in the core circuit (Figure 2(b)).

To model synaptic and gap-junctional precision, we re-fit the model to 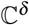 and 𝔾^*δ*^ (Methods). To control for synaptic variability due to differences in axon placement, we restricted our analysis to 𝔸^4^ contact sites (a more general treatment is given in Supplementary Results). For both connection types, our models predict high precision combined with basal connectivity are required to account for the reproducibility and variability across datasets (Figure 2(a)). For the bilateral worm, the synaptic precision of 91% implies a much higher (∼99%) probability of at least one connection (on the left, right or both sides). These results suggest that each dataset can be divided into a common, precisely targeted core circuit and a variable component, and that it should be possible to distinguish between them. Our model further predicts that > 66 % of 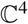 and 𝔾^4^ are good representatives of the core circuit, lending further confidence to the usefulness of the reference connectome. Using model-generated surrogate datasets (Methods), we further find that, on average, each surrogate dataset includes approximately 75% target axon contacts and 25% off-target contacts (Figure 2(b), Figure S6(a)). Thus, we predict that our reference set of axonal contacts is, by-and-large, highly conserved. In contrast to axonal contacts, only 61% of synaptic contacts and 58% of gap junctions form the core wiring in *C. elegans* (Figure S9). In each animal, the conserved core connectome is supplemented by a large set of variable synapses. These results hint at considerably more individual variability and redundancy in wiring than previously expected. To provide developmental context, both axonal and synaptic contacts between early growing nerve ring pioneer neurons^17^ are more reproducible than later growing axons (Figure S10).

To determine the organizational principles supporting this connectome, we now integrate our reference connectome with topographic features across different spatial scales.

### The multiscale connectomes of the *C. elegans* nerve ring coordinate synaptic pathways

The high reproducibility of axon-axon contacts, especially large and early pioneer axon contacts, points to an underpinning developmental program that imposes a high-level spatial organization. To visualize the axonal organization of the nerve ring, we applied a force-directed algorithm to the 𝔸^4^ graph, whereby cells with larger contact areas are placed closer together (Figure S11(a)). Immediately, we see a clustering of neurons that mirrors the anterior-posterior (AP) placement of axons in the nerve ring. Papillary sensory neurons innervate the nerve ring from the anterior while amphid sensory neurons innervate the posterior nerve ring. Both anterior and posterior sensory cells converge onto inter- and motoneurons from both sides. Furthermore, the sublateral pioneer axons (first to innervate the nerve ring^18^) are all placed centrally along the AP axis (Figure S11(a)), strongly suggesting that the nerve ring develops over time from the inside out.

To better identify clusters or subgroups of axons, we used a multi-level graph clustering algorithm (Methods). We find that 5 axon subgroups emerge from the data that are spatially ordered along the AP axis of the worm (Figure 1(a-b)). These subgroups form stereotyped bundles of axons that could structurally support the highly stereotyped axonal trajectories at the single neuron level. All bundles are clearly identifiable both by behavioral functions and anatomical organization (Supplementary Results). We label the bundles *taxis*, *avoidance*, *sublateral*, *lateral* and *anterior*. Based on their neuronal identities and placement on our force diagram, we conjecture that the taxis bundle modulates a number of attractant behaviors to control navigation and foraging; the avoidance bundle integrates nociceptive and mechanosensory cues; the anterior bundle integrates papillary sensory inputs to rapidly control head movements and feeding; the sublateral bundle encodes O_2_/CO_2_ cues; and the lateral bundles integrates nose and body mechanosensory information. The lateral and sublateral bundles also perform key nerve ring integration functions to drive whole body behaviors. Collectively, these results demonstrate an important organizing principle: the nerve ring is partitioned into functional axon bundles ordered along the AP axis, a structure that must be developmentally orchestrated.

By superimposing synaptic sites on the bundle structure (Figure 1(c), Figure S11(b)), we find distinct bands of synapses grossly overlapping the axon bundle organization, and arranged in the same anterior-posterior order (Figure 1(e)). Radially, lamina form concentric rings around the pharynx (Figure 1(f)). Consistent with the spatial adjacency layout (Figure 1(a)), distinct sensory pathways are located in the outer anterior and posterior cylindrical lamina and converge from apposing sides of the nerve ring onto the same interneuron-to-motoneuron pathway (Figure 1(e,f) and S12). Lamina are less apparent for gap junctions (Figure S12(b)) possibly due to their lower density (Table S1). We conclude that the axonal bundle and synaptic laminar organization of the nerve ring achieves both compactness and effectiveness, compressing its distinct functional circuits into an extremely small volume. The laminar pattern of synapses supports the notion of a modular connectome consisting of distinct synaptic pathways. Indeed, the AP and radial organization of synapses could support a hierarchical network topology^5,12,15,16,19^ to achieve sensorimotor convergence, possibly resembling the high-level functional organization reported in vertebrates^20^.

By their very nature, axons connect distinct anatomical regions of the nervous system. While most take an efficient path between one brain region and another^21^, recent reports of three claustrum neurons wrapping around the circumference of the mouse brain^22,23^ point to important exceptions to this rule, and have been suggested to underpin sensory integration roles and brain-wide coordination^24^. In the *C. elegans* nerve ring, we find that ∼40% of axon-axon contacts are made between different bundles (Figure 1(b)). Asking whether designated axons may synaptically couple different bundles, we classified cell classes by their axonal contact with neighboring bundles (Methods, Figure S13, Supplementary Results) and identified 39 single-bundle,^11^ 25 boundary and 30 trans-bundle cell classes. Single-bundle axons are typically located in the middle of their axon bundle and make minimal contact with other bundles (Figure 3(a)). Boundary axons are typically placed at the interface of axon bundles (Figure 3(b)). Trans-bundle axons have spatially distinct axon segments that traverse different bundles (Figure 3(c)).

**Figure 3.**
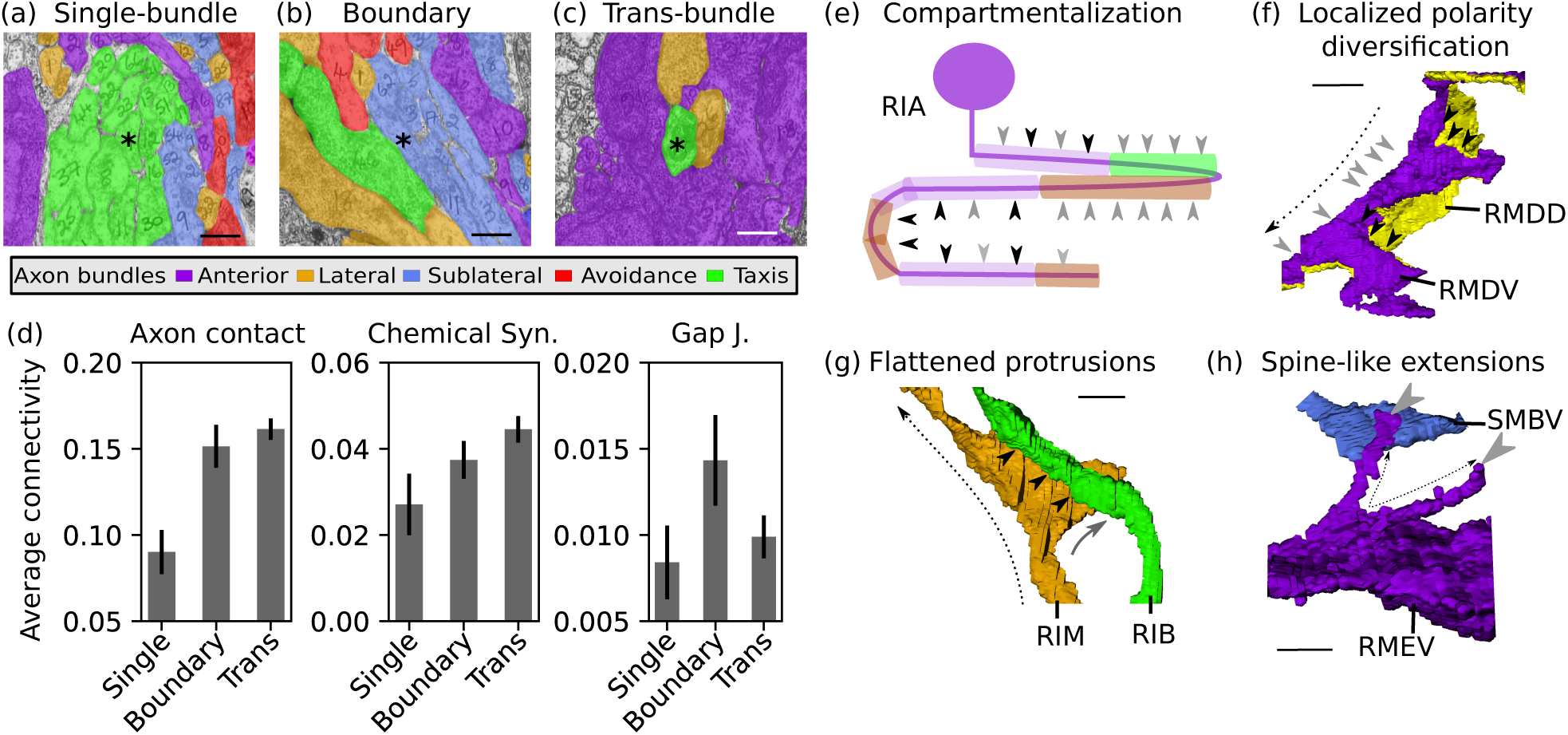
Nano-, micro- and meso-scale axonal structures and organization supports intra- and inter-bundle connectivity. 33 cell classes (6 single-bundle, 7 boundary and 20 trans-bundle) exhibit subcellular organization patterns supporting synaptic connections across bundles. Representative EM sections (scale bar 0.5 *µ*m): (a) Single-bundle axons (*, 39 of 94 cell classes) typically make minimal contact with other bundles. (b) Boundary axons (*,25 cell classes) are placed at bundle interfaces. (c) Trans-bundle axons (*, 30 cell classes) extend outside of their defined bundles. (d) Average connectivity of axon contacts, chemical synapses and gap junctions of the 3 axon types. Connectivity: fraction of total cells contacted by an axon. Error bars: standard error. (e) Schematic of trans-bundle cell RIA demonstrates synaptic compartmentalization. RIA synapses change polarity with changes of neighborhoods. Gray arrows: synaptic inputs. Black arrows: synaptic outputs. Bundle neighborhoods: Anterior (purple), taxis (green) and lateral (brown). See Figure S14(a,b) for volumetric rendering. Volumetric rendering of (f) RMDV and RMDD and (g) RIM and RIB axons, show flattened protrusions that support localized, reproducible synaptic connections. RMDV (f) axons receive synaptic inputs (grey arrows) along their main axonal trajectories (dashed arrows) and extend flattened protrusions where they synapse onto RMDD, diversifying synaptic polarity. RIM axons (g) extend similar protrusions from their main trajectory (dashed arrow, lacking synapses) to synapse onto RIB (black arrows). See Figure S15(a,b) for volumetric rendering. (h) Spine-like extensions (dashed black arrows) grow out of the RMEV cell body to receive synaptic input (grey arrow) from SMBVL and SMBVR. For visual clarity, only SMBVR is shown. See Figure S16(b). (e,f,g,h) All examples are observed on both right and left sides of both the L4 and the adult animals (Figure S2, additional examples in SOM). (e,f,g) Cell colors denote bundle assignment. (h) RMDD (anterior bundle) was colored yellow to visually distinguish it from RMDV. Scale bar: 1*µ*m.

We note that 7 boundary and trans-bundle axons have been identified as the so-called “rich club” *C. elegans* neural circuit (Supplementary Results), a set of neurons that act as circuit hubs through connectivity with each other (high assortativity) and with other neurons^25^. Consistent with this, we find that boundary and trans-bundle axons carry a larger relative load of both axonal and synaptic connectivity in the nerve ring (Figure 3(d)). Thus, we hypothesize that boundary and trans-bundle axons synaptically link spatially distinct parts of the nerve ring to support brain-wide coordinated activity^11^.

To synaptically link distinct regions of the nerve ring, axons must not only reach the designated anatomical region, but synapse appropriately. In *C. elegans*, AIB neurons transition from being pre- to postsynaptic as their axons change neighborhoods at the dorsal midline^11^ (Figure S14). We identified 33 cell classes (20 trans-bundle, 7 boundary and 6 single-bundle) whose axons similarly synapse across distinct regions of the nerve ring (Methods), using two principal strategies: synapse compartmentalization (19/33 cell classes), where different segments of the axon either change synaptic polarity or synaptically target a different axon bundle (Figure 3(e)), and flattened protrusions (23/33 cell classes) that increase an axon’s surface area at discrete points along its trajectory (Figure 3(f,g)). Synapse compartmentalization could facilitate both long-range synapse communication (for isopotential cells like AIB) and subcellular encoding of behavior (for compartmentalized cells like RIA^27^). Flattened protrusions either penetrate neighboring bundles (Figure 3(f)) or locally diversify synaptic polarity (Figure 3(g)) to locally combine synaptic inputs and outputs. In some cases, these protrusions suggest primitive attempts at axonal branches (Figure S15) of dendritic spines (Figure 3(h) and S16). We conclude that boundary and trans-bundle axons synaptically exploit conserved nano- and microscale structures and synaptic compartmentalization to link spatially distinct regions of the nerve ring.

### A *C. elegans* brain map

We integrate the knowledge gained to map the architecture of the *C. elegans brain*: The macro-connectome bundle organization (Figure 1(a)) allows us to crudely divide the wiring diagram into 5 modules and study the circuitry in each module. The meso-connectome core-periphery bundle organization supports the classification of axons as single-bundle and multi-bundle neurons (Figure 3(a-c)), allowing us to map the coordination among bundles, facilitated by micro- and nanoscale axonal structures (Figure 3(e-h)). The reference connectome allows us to focus on reliable, likely conserved connections (Figure 2). Finally, classification of neurons as sensory, interneuron and motoneuron allows us to trace sensorimotor pathways within each bundle. By combining these data, we set out to construct a brain map of the *C. elegans* nerve ring. We posit a parsimonious 3-layer architecture with parallel bundles and assign every neuron of the nerve ring into a layer (Methods). To achieve overall feedforward pathways, sensory neurons all occupy the first layer and multi-bundle neurons dominate layer 3 (Methods, Figure 4).

**Figure 4.**
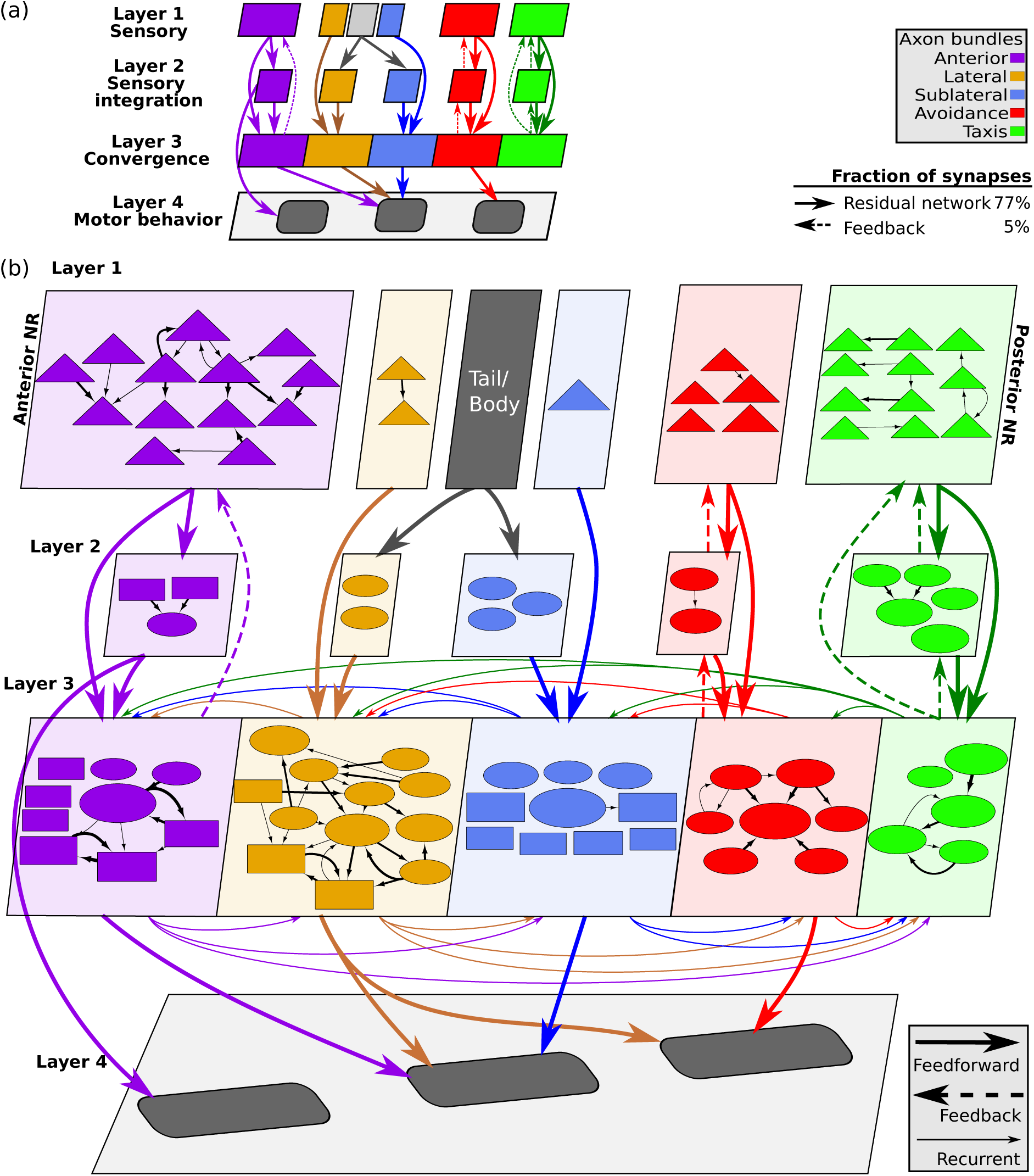
The *C. elegans* brain map. (a) A 4-layer map of synaptic information flow in the nerve ring is well described by a Residual Network architecture^11^. The Residual Network (solid arrows and recurrent connectivity between physically connected blocks in layer 3, spanning 77% of 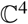 synapses) shows a feed-forward motif for each bundle that converges onto layer 3, supporting functional sensory-motor pathways with strong sensory-motor convergence. Layer 3 consists of primarily trans-bundle interneurons and motoneurons that form a tightly recurrent distributed circuit across all bundles. Dashed arrows: intra-bundle feedback (4%). (b) All 82 bilateral (left/right) homologous neuron pairs and 8 single neurons with reproducible synaptic connectivity in the nerve ring overlaid upon the schematic model in (a). Node sizes are proportional to the synaptic degree (total number of synaptic inputs and outputs). Triangles: sensory neurons. Ovals: interneurons. Rectangles: motoneurons. Cell names have been removed for visual clarity, see Figure S17 for the *C. elegans* brain map with cell names restored.

Connectomic features, identified from network analysis of the *C. elegans* connectome (such as hub and rich club neurons, network motifs and the small world organization as well as new features such as fan-in and fan-out motifs) can now be interpreted within the context of modular, brain-wide computation and information flow. In particular, the feedforward motif, previously identified in the *C. elegans* connectome, reappears in our map as the skeleton of the layered synaptic pathways within each bundle (>80% of all 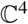 contacts), and gives rise to a Residual Network (ResNet) architecture^28^ (Figure 4(a), SOM, Figures S17-S20 show additional contacts superimposed on the ResNet). Inspired by the layered connectivity of pyramidal neurons in the mammalian cortex, artificial ResNet architectures enhance the resilience of synaptic pathways, and support flexibility and plasticity.

Examination of the *C. elegans* brain map (Figure 4(a)) reveals a number of features: layer 1 separates the bundles (with a few notable and functional exceptions, Figure S17); intra-bundle intra-layer connectivity indicates that sensory neurons likely perform limited sensory computation in addition to sensory encoding of environmental cues, and allows the identification of sensory hub (high-degree) neurons. Layer 2 largely maintains the bundle separation and includes primarily single-bundle neurons, consistent with separate bundles supporting distinct functions. Convergence of sensory neurons onto this sparser layer reveals a fan-in architecture, supporting intra-bundle sensory integration. Layer 3 contrasts with the above. Inputs are received from all three layers: Synapses from layers 1 and 2 are primarily from the same bundle, but layer-3 synapses mix the bundles into a recurrent, highly distributed circuit, consistent with the dominance of boundary- and trans-bundle neurons in this layer and suggestive of brain-wide coordination. Outputs from the nerve ring control the pharynx, head and neck muscles and the motor circuit of the ventral nerve cord (VNC). Pharyngeal output is mediated by layer-2 neurons of the anterior bundle, indicating that the pharyngeal control is independent of the distributed layer-3 circuit. In contrast, head and neck muscles are controlled by three bundles (layer-3 anterior, lateral and sub-lateral bundles) and the VNC is controlled by both lateral and avoidance bundles, revealing the convergence of sensory pathways and associated modular sub-circuits into a small number of highly coordinated motor programs.

## Discussion

The *C. elegans* connectome has been available for over 30 years, and yet the delineation of functions within its main neuropil is still incomplete. By characterizing the spatial embedding of its connectome, we sought insight into the structures that could support a hierarchical, modular and nested architecture in the *C. elegans* brain. We found that the *C. elegans* core connectome is remarkably well described by a modular Residual Network, an architecture noted for its resilience and adaptability. Where previously triplets of neurons were shown to exhibit feedforward motifs^29^, our layerd map reveals a high-level feedforward architectural motif, reminiscent of layered cortical architectures^30^. This ‘connectionist’ description of a biological brain provides a promising methodology for identifying parallel and distributed circuits. Furthermore, as our map is informed by the richness and diversity at the finest scales, it allows for an integrated top-down and bottom-up assignment of cell classes to different pathways and layers.

Superimposed on the Residual-Network template is a hierarchy of recurrent connections, consisting of intra-bundle, intra-layer local computation, akin to local circuits. Within these local circuits, *C. elegans* neurons seem relatively simple: single-bundle axons by-and-large lack structural or functional compartmentalization. Thus, consistent with the neuron doctrine, at the single module level, neurons represent the basic unit of computation. However, the modular architecture converges within the final layer to achieve brain-wide coordination of behavior. In this distributed circuit, the nano-connectome rules: highly specialized subcellular structures give rise to compartmentalized dynamics and inter-link distant regions of the *C. elegans* brain. Similar subcellular structures performing analogous functions, found in thalamic local interneurons^31^, reveal a richness of subcellular computation and hint at common developmental mechanisms.

The core-periphery structure of axon bundles in *C. elegans* is consistent with reproducible axonal placement and trajectories which support the large number of conserved axonal contacts. This mesoscopic spatial arrangement of axons thus supports and constrains the synaptic architecture of the animal: centrally placed axons synapse almost exclusively within their bundle. The collection of single bundle neurons thus sheds light on each bundle’s sensory and sensory-integration functions. Brainwide coordination is achieved downstream of single bundles by designated multi-bundle axons that interface between or thread across multiple bundles, thus underpinning sensory-convergence and sensorimotor transformations. This brain map and its nested architecture might suggest a much closer analogy between the *C. elegans* neuropil and the coordination between the nano- and macro-connectome of other invertebrates and even vertebrates^20^.

The concept of a reference connectome was key to our brain map and the modeling framework we used to establish this reference can easily be extended to accommodate future connectomes. In vertebrates, nanoscale organization underpinning individual synapses is highly variable, supporting individual wiring, plasticity and adaptability. In *C. elegans*, the proportion of conserved synapses was unknown. We found that the connectome consists of a core, conserved circuit that is embedded in a significant variable background. Our identification of a candidate core connectome set the bar for a multi-scale topographic organization of the nerve ring. If the core circuit represents the baseline functionality of the animal, the variable component could support redundancy, individuality and plasticity. If so, it would seem that both circuits must be spatially constrained to ensure their functionality.

Despite detailed characterization of neuronal morphologies and axonal trajectories, the *C. elegans* nerve ring is typically viewed as a disorganized tract of axons, limiting the value of structure-function analysis of the connectome. However, the large number of cell classes, so densely packed in the nerve ring, present a challenge to physically achieving stereotyped connectivity. Our finding of finely orchestrated organization across scales imposes spatial constraints on axon and synaptic placement, thus restricting each neuron’s connectivity problem to a local neighborhood. This scalable solution naturally generalizes to much larger nervous systems. Viewed differently, the spatial organization reduces the required capacity for cell-cell molecular recognition machinery, while increasing the complexity of mechanisms producing the cell’s morphology and relative positioning in the tissue. But how is the bundle organization developmentally orchestrated? Previous models of neuropil development have proposed that pioneer axons guide follower axons^32^. While such models could be generalized to identify the pioneer axon of each bundle, the fine core-periphery (single-boundary axon) bundle structure indicates a more elaborate developmental mechanism. In one such model, some guidance molecules would coordinate the relative bundle placement and others – the placement of axons within bundles. Identifying key guidance molecules in early nerve ring formation (e.g. SAX-3/Robo^17^) may help to address such predictions. Whatever the developmental mechanisms may be, the brain map of *C. elegans* requires that these mechanisms too are nested and coordinated across scales to guide and support the modular, scalable and flexible neural architecture that produces the mind and behavior of the nematode *C. elegans*.

## Methods

See Supplementary Methods.

## Acknowledgments

We thank Jonathan Hodgkin and John White for their help in donating archival TEM material from LMB/MRC to the Hall lab for curation. CB was supported by the Leeds International Research Scholarship. DHH is supported by NIH OD 010943. NC was supported by EPSRC EP/J004057/1.

## Author contributions

CB, SC and SE conceived the project. CB and SC segmented the electron micrographs. CB and NC analyzed data. CB and NC wrote the manuscript. DHH curated the data. SC, DHH and SE provided critical revisions.

